# Metagenomes and Metagenome-Assembled Genomes from Microbial Communities in a Biological Nutrient Removal Plant Operated at Hamptons Road Sanitation District (HRSD) with High and Low Dissolved Oxygen Conditions

**DOI:** 10.1101/2025.11.10.687637

**Authors:** Blaise M. Enuh, Kevin S. Myers, Charles Bott, Stephanie Klaus, Kester McCullough, Lilian McIntosh, Natalie Beach, Michelle Young, Timothy J. Donohue, Daniel R. Noguera

**Author notes:** Corresponding author: Daniel R. Noguera.

## Abstract

Aeration in biological nutrient removal (BNR) systems constitutes one of the largest energy demands in water resource recovery facilities (WRRFs). Previous studies have shown that lowering dissolved oxygen (DO) concentrations can sustain effective nitrification and phosphorus removal while substantially reducing energy consumption. However, the microbial mechanisms enabling these low-DO processes remain poorly understood. In this study, we investigated microbial communities associated with reduced DO in a pilot-scale BNR system operated by the Hampton Roads Sanitation District (HRSD). DO was reduced over an 18-month period from 2.5 mg/L to 0.2 mg/L. Metagenomic DNA was obtained from samples from each DO condition then sequenced using PacBio HiFi technology. A total of 316 metagenome-assembled genomes were recovered and after dereplication, 207 were found to be unique. These data augment the metagenomic information related to wastewater treatment under low-DO conditions and provide valuable resources for understanding microbial adaptation to oxygen-limited BNR operation.

## Background

Aeration within biological nutrient removal (BNR) systems can account for a large fraction of the electricity costs at water resource recovery facilities (WRRF). Several studies have demonstrated that by reducing dissolved oxygen (DO) levels in BNR processes, it is possible to maintain efficient nitrification and phosphorus removal with lower energy consumption (Fitzgerald et al., 2015; Keene et al., 2017; Stewart et al., 2021). Lowering DO in aerated sections of BNR plants leads to an adaptation of the microbial community to the new environmental conditions (Fitzgerald et al., 2015; Park and Noguera, 2008, 2004). The underlying dynamics of these microbial community changes when DO is reduced remain poorly understood.

Here, we report on the metagenomes and metagenome-assembled genomes (MAGs) from a pilot plant operated by the Hamptons Road Sanitation District (HRSD) when the DO was reduced in the aerated portions of the treatment train. Activated sludge samples were collected from the pilot plant at the beginning of the operation when the target DO was high (2.5 mg/L), and at the end of the operation, 18 months after the target DO was lowered (0.2 mg/L). In total, 4 samples were analyzed, 2 obtained during high-DO operation and the other 2 from low-DO operation.

### Metagenomic DNA extraction, library preparation, sequencing, and analysis

DNA was extracted using the DNeasy PowerSoil Kit using the published protocol (Qiagen, Germantown, MD), quantified using a Qubit fluorometer (Fisher Scientific, Waltham, MA), and stored at -20°C until sequencing. The quality of the extracted DNA was measured on a NanoDrop One (Fisher Scientific). Quantification of the extracted DNA was done using the Qubit dsDNA High Sensitivity kit following standard protocols (Fisher Scientific).

The samples were sent to the University of Wisconsin-Madison Biotechnology Center (Madison, WI) for library preparation and sequencing. A HiFi library was prepared according to PN 102-166-600 Version 04 workflow (Pacific Biosciences, Menlo Park, CA) with standard parameters. Briefly, the procedure involved fragmentation with g-TUBEs (Covaris, Woburn, MA) according to the shearing protocol for large genomes (2,164 x g) and size selection using diluted AMPurePB beads (Pacific Biosciences) as detailed in the procedure-checklist. Library integrity was evaluated on the FemtoPulse System (Agilent, Santa Clara, CA). The library was quantified using the Qubit dsDNA High Sensitivity kit and sequenced on a Sequel II using Sequel Polymerase Binding Kit 2.2 following the standard protocol (Pacific Biosciences). Default parameters for all programs were used unless otherwise stated. Separate sequencing runs were done at each sampling period for the two DO end points. The Circular Consensus Sequence (CCS) reads were assembled utilizing either metaFLYE (v2.9-b1768) (Kolmogorov et al., 2020) polished with racon (v1.4.20) (Vaser et al., 2017) for the high-DO samples, or metaMDBG (v0.3) (Benoit et al., 2024) with a racon module (v1.4.20) (Vaser et al., 2017) for the low-DO samples. Subsequently, the reads were mapped onto the assemblies using minimap2 (v2.22-r1101) (Li, 2018). Binning of the assemblies was conducted with metaBAT2 (v2:2.15) (Kang et al., 2019). Identification of contaminated contigs within each bin was done with ProDeGe (v2.3) (Tennessen et al., 2016) and with custom scripts used for tetranucleotide frequency analysis (run.GC.sh, Calculating_TF_Correlations.R; https://github.com/GLBRC/metagenome_analysis). Contigs identified as contaminated through both methods were removed from the final MAG assemblies. All MAGs were evaluated for quality using CheckM (v1.2.2) (Parks et al., 2015) and the taxonomy of each MAG was determined using GTDB-Tk (v2.1.0) database release 09-RS214 (Chaumeil et al., 2020). Functional annotations for each MAG were assigned using Bakta (v1.9.1) (Schwengers et al., 2021).

RAxML-NG adaptive (v1.2.1) (Kozlov et al., 2019) with default settings was used to infer the best phylogenetic tree by maximum likelihood estimation based on the multiple sequence alignment of 120 bacterial marker genes generated by GTDB-Tk. The resulting tree was visualized in TreeViewer (v2.2.0) (Bianchini and Sánchez-Baracaldo, 2024) and annotated in Inkscape (v1.2.2) (The Inkscape team, 2025). The finalized tree is shown in Figure 1.

**Figure 1.**
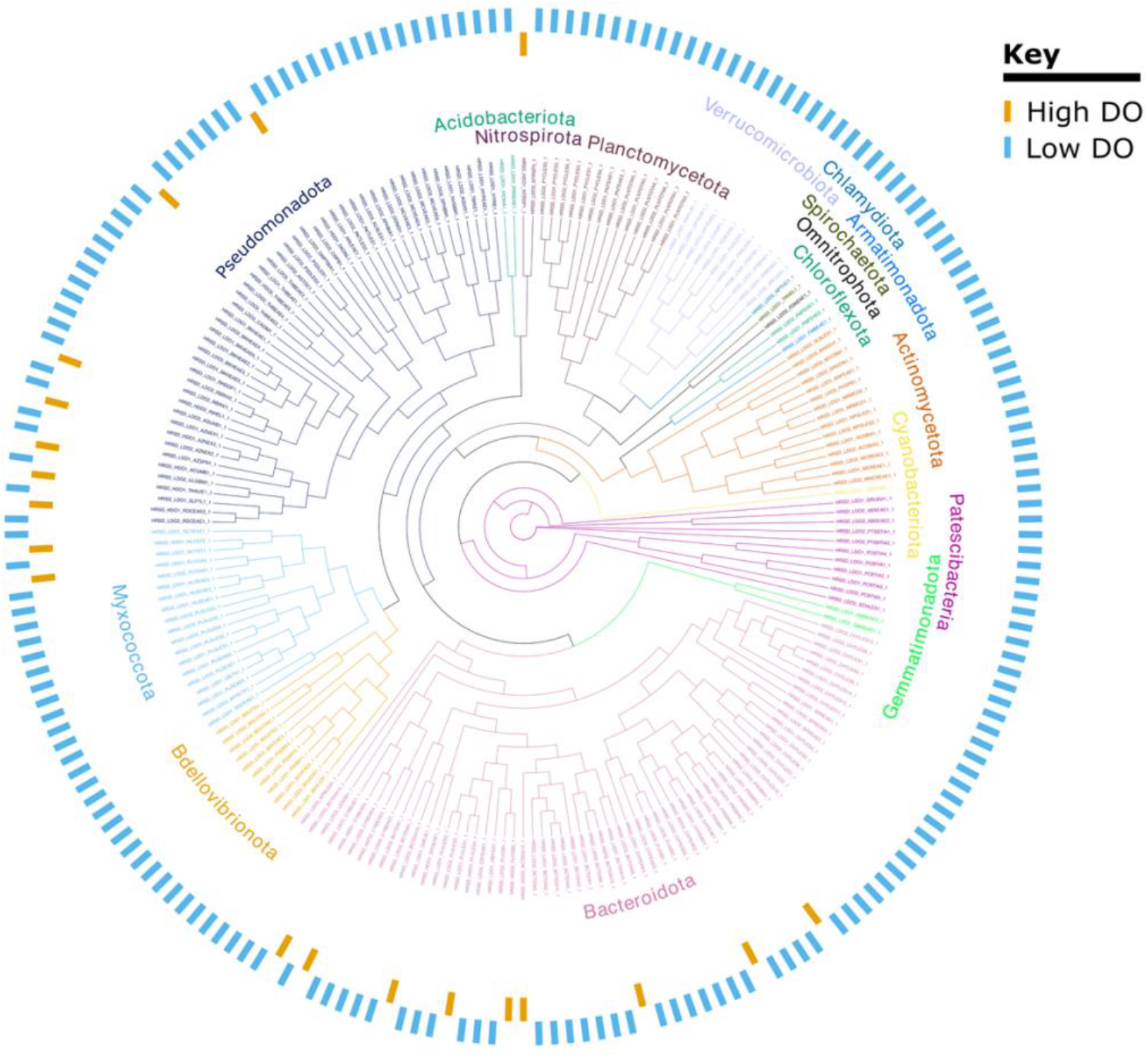
Phylogenetic map showing an overview of community structure based on the 207 unique MAGs from the HRSD plant assembled from both low and high DO samples. From the innermost to the outermost ring: The innermost ring represents phylogenetic clustering of MAGs. The names of the phylum are colored according to the clusters they represent. The next ring in orange points to MAGs that were assembled from the high DO samples while the blue ring points to MAGs that were assembled from the low DO samples.

In total, we obtained 316 MAGs from the 4 samples (Supplementary File S1), with 278 MAGs from the low DO samples and 38 from the high DO samples. These MAGs were dereplicated with dRep (v0.6.1) (Olm et al., 2017), resulting in a set of 207 unique MAGs with over 75% completeness and less than 10% contamination (Figure 1). These data augment the metagenomic information related to wastewater treatment under low-DO conditions. The MAGs obtained and their characteristics are provided in the Supplementary File S1.

## Acknowledgments

This work was supported by funding from the U.S. Department of Energy (DOE), Office of Energy Efficiency & Renewable Energy, under award DE-EE0009509, and partially based upon work at the Great Lakes Bioenergy Research Center supported by the U.S. Department of Energy, Office of Science, Biological and Environmental Research under Award Number DE-SC0018409. DNA sequencing was done at the UW Biotechnology Center’s DNA Sequencing Facility (Research Resource Identifier – RRID:SCR_017759).

## Data availability

Fastq files for the 4 metagenomes have been deposited in the NCBI database under Bio project number PRJNA1156690. The MAGs and their annotation dataset can be accessed on Figshare (10.6084/m9.figshare.30389092). The custom scripts used in these analyses are available at the GLBRC GitHub repository (https://github.com/GLBRC/metagenome_analysis).

